# Reproducible transversal mouse brain sections using low-cost 3D printable resin matrix

**DOI:** 10.64898/2026.07.27.741057

**Authors:** Kelly Falcón, Arnaldo Bisbal López, Rita Thammakhoune, Hope Ayim, Matthew Clark Jung, Anish Krishna, Camila C. Aragón, Aidan Conley Kieffer, Tuan Leng Tay

## Abstract

Rodent brain matrices that produce coronal or sagittal brain sections for histology confer reproducibility and enable high throughput processing of tissues. However, a stainless steel or acrylic brain matrix that produces tissue sections in a transversal (or horizontal) orientation is currently unavailable as a standard tool. This limits the direct comparison of bilateral brain hemispheres within a single histological section, as freehand trimming to obtain horizontal planes is not easily replicable across samples. To mitigate this challenge, we designed a low-cost (USD 7 per unit), 3D-printed resin-based transverse brain matrix that accommodates mouse brains ranging from 12 to 16 mm in length from the olfactory bulb to the brainstem. Our matrix reproducibly generates horizontal tissue sections with a minimum of 1-mm-thickness without causing visible tissue deformation, which is comparable to the performance of commercial rodent brain matrices. Users may adapt the accompanying CAD code using our video tutorials to customize the transverse brain matrix for their specific needs, including alternative brain size, shape, and tissue thickness.

## INTRODUCTION

Histological analysis is a gold standard for studying neuroanatomy and brain cytoarchitecture *in situ*. Guided sectioning that yields robust and reproducible brain tissue samples is achieved using brain matrices. Efforts to standardize the brain sectioning workflow have led to the development of multiple brain matrices constructed using computer-aided design (CAD) and additive manufacturing better known as 3D printing technology (Holtz et al., 2026; Huang et al., 2019; Lee et al., 2022; Tyson et al., 2015; Yamazaki et al., 2025). Current standard brain matrices are constrained to generate coronal or sagittal planes. As a result, direct investigation of bilateral elements that traverse the anteroposterior and dorsoventral orientations is limited. With the availability of only coronal and sagittal mouse brain matrices, quantitative spatial analysis of neural connectivity and cellular organization in transverse (or horizontal) planes must be performed across multiple fragmented sections (Baccini et al., 2021). Investigators seeking to examine bilateral brain hemispheres have relied on freehand trimming that introduces variability in slice angle and thickness, thereby reducing inter-sample consistency (Defazio et al., 2015; Huang et al., 2019; Lee et al., 2022). These technical limitations severely hinder the study of interhemispheric cortical communication through the thalamic commissures (Szczupak et al., 2021), anterior commissural projections involved in avoidance and appetitive behaviors (Tian et al., 2024), and bilateral epileptiform activities in the hippocampal commissure (Liu et al., 2021), among other examples. Horizontal brain sections preserve the entire length of a white matter tract within a single histological plane, which would offer a unique advantage for studying spatially non-uniform pathological processes. For example, coronal sections prevent visualization of the origin and full trajectory of the developing corpus callosum (cc) axons and their association with microglia, the brain-resident macrophages required for axonal growth and guidance (Pont-Lezica et al., 2014). Studying how the white matter tract matures into a heavily myelinated structure with the tightly regulated production of myelin basic protein and myelin proteolipid protein in the adult brain and knowing how to prevent dysfunctional release of these proteins, are critical to our fundamental understanding of neurological diseases such as multiple sclerosis and Alzheimer’s disease (AD) (Ruskamo et al., 2022; Sobue et al., 2023). In humans, studying the topography of white matter fibers in the midsagittal cc from whole-brain reconstruction of coronal sections has been challenging due to discontinuous sampling in whole-brain tractography (Zhang et al., 2023). To date, no preclinical study of mammalian cc along the rostrocaudal rodent brain has been done despite the known vulnerability of the cc in AD pathogenesis, indicating a critical and urgent methodological gap that could be addressed by having the appropriate research tools. While high-resolution horizontal mouse brain atlases derived from high-throughput micro-optical sectioning tomography (MOST) using brains embedded in plastic resin have been reported (Feng et al., 2025; Li et al., 2010), the advanced MOST technology is not available to most researchers who work with fresh frozen or fixed frozen brain tissues. Horizontal rodent brain sections have reportedly been cut from low-cost gelatin-embedded preparations that can be scaled (Nwafor et al., 2024), but cryoprotection and embedding procedures take several days and are incompatible with rapid processing of small batches of samples that is typically achieved in a few hours using standard coronal and sagittal brain matrices.

Here, we designed and fabricated a transversal murine brain matrix for reproducible blocking and sectioning along the horizontal plane of an adult mouse brain. Our high-resolution resin-printed device incorporates geometric features to maintain consistent alignment of the whole brain and reduce variability associated with manual freehand trimming. Integrated clamping surfaces distribute pressure across the tissue to reduce deformation during cutting. Reinforced cutting guides were integrated to ensure smooth and repeatable cutting trajectories that generate transverse brain tissue sections with sub-millimeter precision. Comparison of neuroanatomy and nuclei density in mouse brain cryosections acquired with our transversal matrix by multiple independent users against results from coronal and sagittal preparations using commercial matrices demonstrates the simplicity and reproducibility of our new resin-based device.

## MATERIALS AND METHODS

### Brain Matrix Design and Fabrication

The transverse mouse brain matrix was modeled using Onshape (version 1.211, https://bu.onshape.com/). To establish the base of the matrix, a 25.4 mm by 31.75 mm rectangle was drawn using the Sketch and Center Point Rectangle tool and refined using the Dimension tool. The rectangle was extruded by 25.4 mm in the positive z-axis direction using the Extrude tool to form the 3D shape of the matrix. Next, a 14.986 mm by 6.096 mm ellipse was drawn in the center of the device using the Sketch and the Center Point Rectangle tool and finetuned with the Dimension tool to situate the brain tissue chamber. The chamber was designed to accommodate a range of sizes of whole brain harvested from mice at 6 weeks to beyond 24 months old (Allen Institute, 2004). The Line tool was utilized to divide the ellipse into two symmetrical halves. To create the cavity, the Revolve tool was used to rotate the top half of the ellipse 90° towards the negative z-axis and produce a curved concave chamber that conforms to the dorsoventral profile of the brain. The curvature cradles the brain by spreading its contact across the tissue surface and reducing localized compression, while keeping the specimen consistently positioned during blocking and sectioning. One face of the cavity was extruded laterally by 4.191 mm to fit the dorsal-ventral axis of an adult mouse brain. Next, the cavity was extruded by 6.350 mm using the Extrude tool to accommodate the horizontal width of the brain. Starting from the cavity wall that would be in contact with the ventral part of the brain, the Line tool was used to draw a 0.559-mm-thick line that would eventually form the ventral-most slit of the brain matrix to contain a blade that will hold (or block) the brain in place. To obtain 1-mm horizontal brain sections, the Line tool was used to create a 1.003-mm wedge that alternates with the slit. The 0.559-mm slit was extruded 11.430 mm in the negative z-axis to establish the depth of the slit. This extruded slit was replicated using the Linear Pattern tool towards the opposite cavity wall (dorsal) to achieve an instance count of five and ensure a 1.562-mm distance between two slits. Diagrammatic representation of the CAD process is provided in the **Supplementary Methods** and the code for 3D printing is publicly available (Falcón and Tay, 2026). Brain matrices were printed using a Form 3+ Resin 3D Printer (Formlabs, Boston, USA) and the PreForm software (version 3.55.1, Formlabs) at the Boston University (BU) Engineering Product Innovation Center (EPIC). Gray Resin V4 (#RS-F2-GPGR-04, Formlabs) was used as the printing material.

### Mouse Brains

Juvenile and adult C57Bl6 mice of both sexes were euthanized by carbon dioxide asphyxiation. These animals, which were bred in the BU Animal Science Center with *ad libitum* access to food and water under a standard 12-hour light / 12-hour dark schedule, were no longer used in the authorized research projects. In keeping with the 3R principle of Replace, Reduce, and Refine, we utilized these excess anonymized mice to test the transversal brain matrix prototypes and perform validation experiments. Due to the required anonymization, the phenotypes and precise ages of the animals are unavailable. After decapitation, the mouse brains were carefully removed from the skulls and fixed for 24 hours at 4°C in 4% paraformaldehyde dissolved in 1X phosphate-buffered saline (PBS) solution adjusted to pH 7.4. Fixed brains were rinsed 5 times for 5 minutes each at room temperature in 1X PBS before cryoprotection at 4°C in 1X PBS containing 30% sucrose until the brains were fully immersed. The dimensions of each brain were measured using a digital caliper.

### Brain Sectioning, Embedding, and Slicing

Ultra Disposable Microtome Blades (3053835, Epredia™) were used to block and section the cryoprotected mouse brains in the transverse, coronal (stainless steel, 69-2175-1, AgnTho’s AB), and sagittal (acrylic, 68-1275-1, AgnTho’s AB) brain matrices. Any brand of single-cutting-edge blades with 0.56-mm thickness or less can be used with our transverse resin matrix. After positioning the fixed mouse brain in the cavity of the transverse brain matrix, the brain can be sectioned by several approaches: (1) simultaneous cutting with five blades in all five slits to obtain four 1-mm sections; (2) blocking the dorsal- or ventral-most slit, cutting the next consecutive section without removing the blade to block for the next section, then using the first blocking blade to cut the next section, repeating until all required sections are obtained; (3) blocking with one blade in the dorsal-most (#1) and/or ventral-most slit (#5) and acquiring the required tissue thickness using the remaining slits. The horizontal tissue block between slits #2 and #4 was used for histological analysis. Coronal tissue blocks were obtained between slits #7 (rostral) and #11 (caudal). Sagittal tissue blocks of each hemisphere were obtained between slit #2 (lateralmost) and the medial-most slit. Each brain block obtained from a matrix was immediately embedded in a Tissue-Tek^®^ Cryomold measuring 15 mm × 15 mm × 5 mm (NC9464347, Andwin Scientific) using Tissue-Plus™ O.C.T. Compound (23-730-571, Fisher Scientific) and stored at -80°C. Olfactory bulbs were removed for optimal positioning of the horizontal brain blocks in the cryomold. Tissue blocks were equilibrated to -20°C overnight before slicing at 20-μm thickness using the cryostat (CryoStar NX50, Epredia) and collected on Superfrost™ Plus Microscope Slides (12-550-15, Fisherbrand™). Tissue slices were air-dried and stored at -20°C.

### Hematoxylin & Eosin Staining

Slides were placed in a 24-slide SHURStain stainer rack and well (SS-WLG, General Data Inc) to thaw for 1 minute at room temperature. Tissue slices were rehydrated in 1X PBS for 3 minutes with gentle dipping of the rack for 5 times. Slides were rinsed in the well with cool tap water changed 4–5 times then dipped in Shandon^TM^ Gill^TM^ hematoxylin (6765009, Epredia) for 3 seconds. The blue-stained tissues were rinsed in cool tap water with at least 4 changes until the water was clear without any blue tint. The slides were then dipped in 1X PBS for 20 seconds, followed by a 1-minute-long dip in cool tap water. Next, the slides were incubated stepwise in 70% and 95% molecular biology grade absolute (200 proof) ethanol (BP2818500, Fisher Scientific) for 30 seconds each and counterstained in Shandon^TM^ Eosin-Y stain (6766007, Epredia) for 20 seconds. Tissue slices were dehydrated in 2 changes of 95% ethanol for 15 seconds each, followed by 2 changes of 100% ethanol for 15 seconds each. The slides were subsequently dipped in 3 changes of histological xylene (X3P1GAL, Fisher Scientific) for 1 minute each and left to air-dry under the fume hood for 20 minutes. Slides were coverslipped using limonene mounting medium (ab104141, Abcam) and Superslip cover slip no. 1.5 (12-541-055, Fisher Scientific) and left to cure at room temperature for at least 24 hours.

### Microscopy and image analysis

Brightfield images of horizontal, coronal, and sagittal whole-brain sections were acquired on the ZEISS AxioScan 7 slide scanner using the 20X/0.8 objective lens and high-resolution Axiocam 705 color CMOS camera. The regions of interest (ROI) containing the cc or hippocampus (HIP) were cropped and processed on Zen 3.5 (ZEISS). Measurements and nuclei counts were acquired using standard tools in Zen 3.5 and Fiji (Schindelin et al., 2012). For the horizontal sections, bilateral measurements for 3–9 sets of cc ROI, 9–18 sets of HIP ROI, and 8–18 sets of HIP nuclei in 250 µm × 250 µm ROIs were obtained for each brain. For the coronal sections, bilateral measurements for 4 sets of cc ROI, 4 sets of HIP ROI, and 4–8 sets of HIP nuclei in 250 µm × 250 µm ROIs were obtained for each brain.

### Statistical analysis

Wilcoxon signed-rank test was performed using GraphPad Prism 11 (GraphPad Software, CA, USA). Data are presented as the mean ± standard error of the mean (SEM). *P* < 0.05 is considered statistically significant.

## RESULTS AND DISCUSSION

### Design and application of the transversal brain matrix

Our customized transversal mouse brain sectioning matrix was conceptualized using CAD **(Figure 1 and Supplementary Methods)**. The prototype that produced even and rigid wedges was 3D printed in resin using additive manufacturing (Falcón and Tay, 2026). Our matrix costs ∼US$7 per unit to produce, making it highly cost-effective––relative to commercially available stainless steel or acrylic brain matrices––and accessible to neuroscientists around the world. For researchers without access to an in-house resin printer, our open-source CAD files may be used through institutional makerspaces, university fabrication cores, or commercial 3D-printing services that accept user-submitted CAD or STL files. Three independent testers reproducibly cut horizontal sections using the transversal brain matrix on juvenile and mature adult mouse brains ranging from 12– 14 mm (N = 6) and 14.01–16 mm (N = 6), respectively **(Figure 2A)**. The resin provided sufficient stability and rigidity to the device, ensuring that any tissue placed in the chamber is securely held and that the slits seamlessly accommodate the blades **(Figure 2B)**. In contrast to stiffer materials (e.g., stainless steel and acrylic) used in the production of commercial brain matrices, users of this resin-based device should acquaint themselves with the moderate flexibility of the wedges that separate the slits during initial use **(Figure 2C)**. Cutting angle, blade placement for blocking, and sectioning force and speed may require optimization for individuals to obtain the ideal transverse mouse brain sections **(Figure 2D)**. The Gray Resin V4 used to create our customized transversal matrix may be chilled but cannot be sterilized by autoclaving. The resin-based matrix tolerates ethyl alcohol and can be soaked in warm water with mild detergent to remove dried tissue. Immediate cleaning after use is recommended. The matrix can be gently scrubbed clean under running water using a fine-bristled brush or toothbrush without damaging its surfaces. While material costs, printer resolution, and material compatibility will vary across institutions and geographical locations, we expect minimal barriers for implementation, making our transversal brain matrix a practical and accessible solution to an unmet need in brain histology.

**Figure 1.**
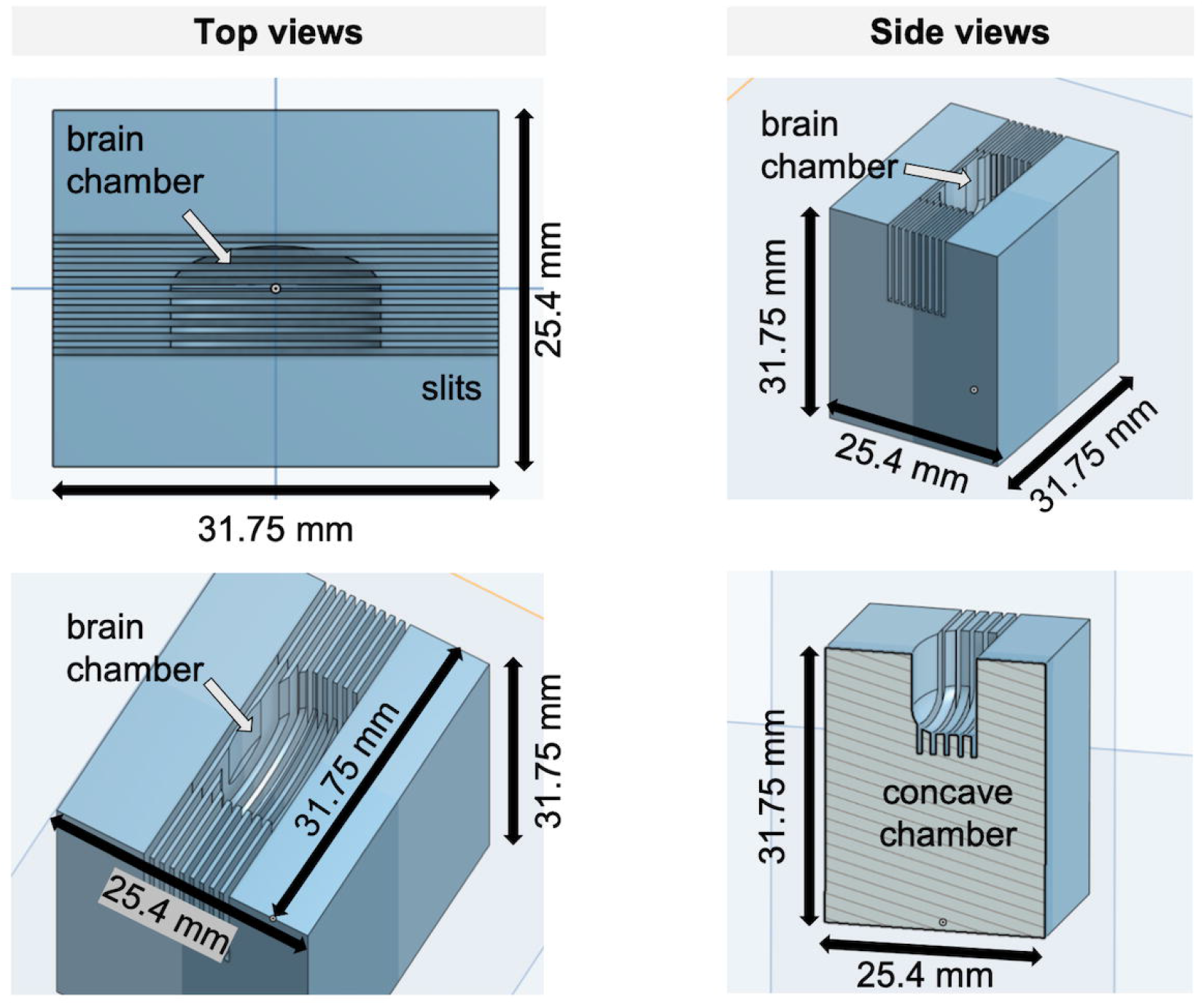
Conceptualization of the transversal mouse brain matrix using CAD to create the concave chamber for holding the brain sideways and slits for placement of the cutting blades. See **Supplementary Methods** and **Supplementary Videos 1–5**.

**Figure 2.**
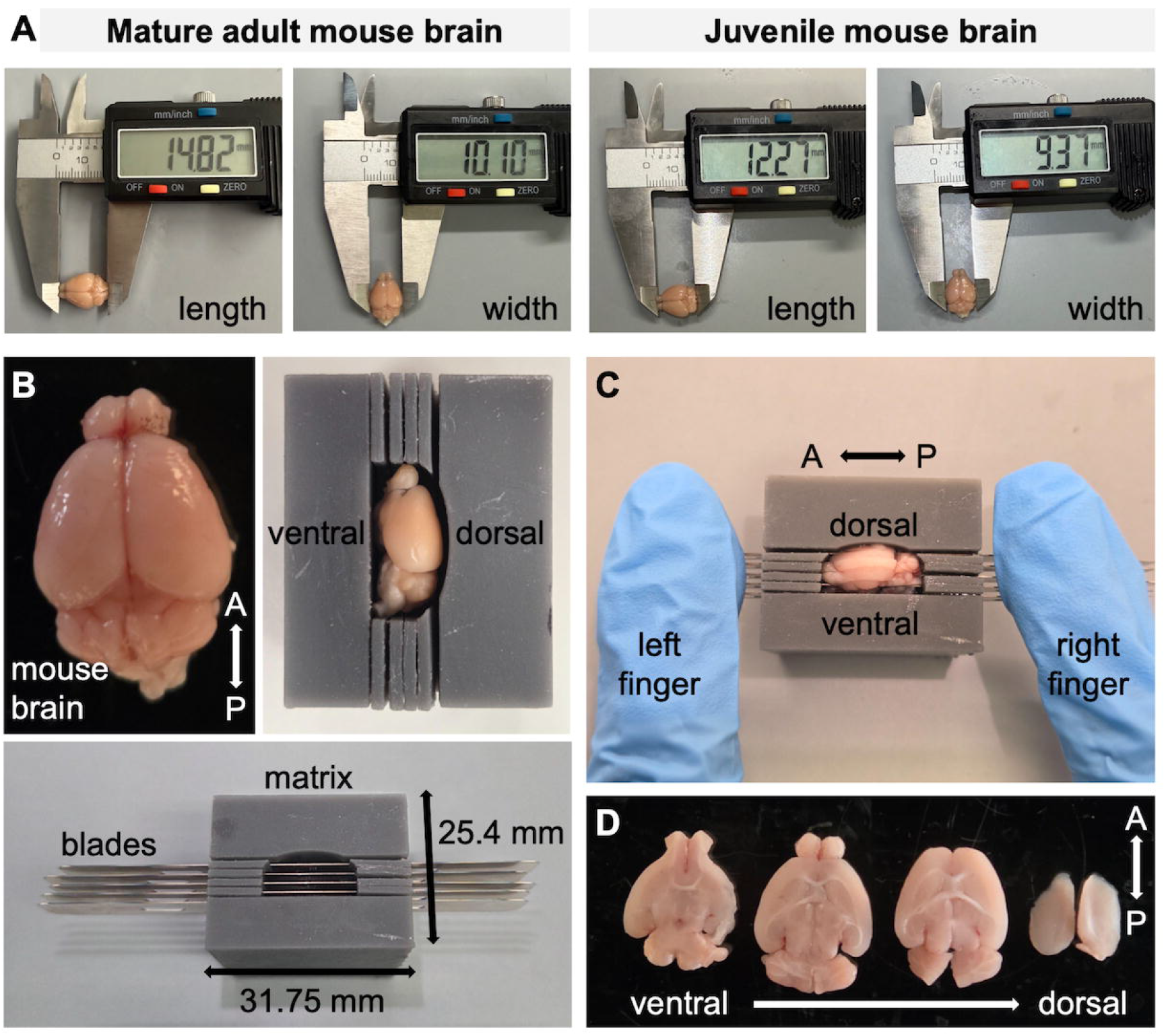
Transverse mouse brain sectioning with the 3D printed resin matrix. **(A)** Classification of test mouse brains based on size: mature adult brains were categorized as 14.01–16 mm in length (N = 6) and juvenile brains were 12–14 mm in length (N = 6) from the olfactory bulb to the brainstem. The widths of all brains ranged from 9.19–10.6 mm. **(B)** Orientation and positioning of an adult mouse brain and cutting blades in the transverse brain matrix. **(C)** Blocking and sectioning of a mouse brain by simultaneous exertion of cutting force using five blades. A swift cutting motion of under 1 sec is recommended. **(D)** Representative consecutive 1-mm sections from simultaneous cutting using five blades. Four brains were cut using this method by 2 independent users. A, anterior; P, posterior.

### Customization of the transversal brain matrix

The design of our brain matrix can be modified to accommodate different experimental requirements, including changes in sectioning angle, brain dimensions, cavity curvature, section geometry, and blade dimensions. To alter the angle of sectioning, select Sketch 2 and use the Transform tool to rotate the ellipse and its bisecting line to the desired orientation **(Supplementary Video 1)**. During this process, select the outer perimeter of the ellipse followed by the bisecting line to ensure proper rotation of the sketch geometry. Following the angular adjustment, edit Extrude 4 to select the entire Sketch 3 and confirm that the sectioning features are extended to span the full width of the modified cavity. To accommodate wider rodent brains, the depth parameter of Extrude 3 can be increased to generate a deeper cavity **(Supplementary Video 2)**. To accommodate larger brains in the dorsoventral axis, the depth of Extrude 2 can be increased to expand the cavity height. If the sectioning slits no longer intersect the cavity following this adjustment, the remaining faces of Sketch 3 should be added to Extrude 4 to restore continuity of the slits **(Supplementary Video 2)**. For longer brains, the major axis dimension of the ellipse in Sketch 2 can be modified by selecting the existing dimension and entering the desired value **(Supplementary Video 3)**. Cavity curvature can be adjusted by changing the minor axis dimension of the ellipse in Sketch 2. Substantial increases in curvature may require corresponding increases in slit depth and, in some cases, an increase in the total number of sectioning slits to ensure complete coverage of the cavity. Section geometry can be modified by adjusting the depth of Extrude 4 to increase or decrease slit depth **(Supplementary Video 4)**.

The number of sectioning slits can be altered by editing Linear Pattern 1 and modifying the instance count parameter. Slit spacing can be adjusted by increasing or decreasing the distance parameter within Linear Pattern 1. Note that because this value represents the combined thickness of the blade and desired tissue section, changes to this parameter will directly affect the thickness of the resulting brain sections when blade thickness remains constant. Modifications to slit spacing requires corresponding adjustments to the instance count to maintain adequate cavity coverage. If cavity height exceeds the range covered by the patterned slits, the Second Direction option within Linear Pattern 1 can be enabled and the left edge of Extrude 3 is selected as the secondary pattern direction. Additional instances can thereafter be generated along this direction by modifying the second-direction instance count to extend slit coverage to the ventral part of the cavity. The distance in both directions must remain the same. To accommodate alternative blade dimensions, the width of the lowermost rectangle in Sketch 3 can be modified to match the desired blade thickness **(Supplementary Video 5)**. The default slit width is 0.559 mm (0.022 in). Accommodation of taller blades can be achieved by increasing the slit depth through modification of Extrude 4 as described above.

### Reproducibility of transversal brain matrix is comparable to standard matrices

To validate the performance and reproducibility of our transversal brain matrix, three independent users were assigned to cut 2–3-mm-thick horizontal (N = 12), coronal (N = 4), and sagittal blocks (N = 4) from cryoprotected whole adult brains and embed them in TissueTek before subsequent slicing on the cryostat. The blade placements were executed respectively in the new transversal matrix and commercially available standard coronal and sagittal matrices to show three different perspectives of the neuroanatomy corresponding to the HIP area seen in the horizontal tissue slices **(Figure 3)**. The resulting tissue sections were randomized for data collection. Quantification of bilateral white matter tract thickness, length across the HIP region, and number of nuclei within corresponding ROIs showed no difference between both hemispheres when the transversal and coronal matrices were used **(Figures 3A and B)**. Furthermore, each orthogonal view reveals a different perspective of the brain compartments and neural circuitry **(Figures 3A–C)** (Franklin and Paxinos, 2008; Rosen et al., 2000), indicating a versatile and advantageous application of our transversal brain matrix as a new standard tool to study bilateral brain nuclei and whole-brain connectivity between intact brain hemispheres.

**Figure 3.**
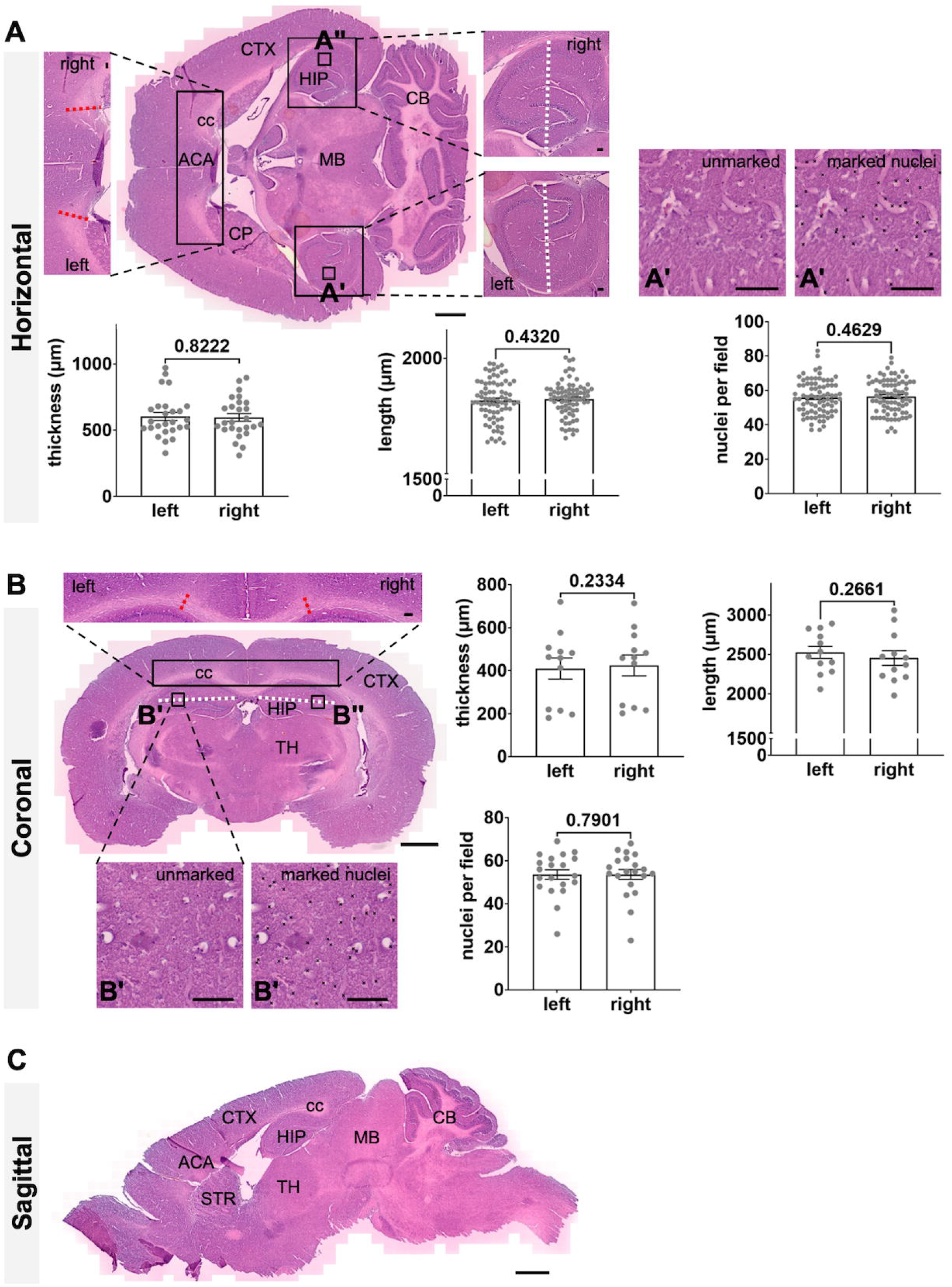
Transversal mouse brain matrix produces 2.5-mm-thick horizontal blocks to obtain 20-µm-thick fixed frozen sections for hematoxylin and eosin staining with reproducible bilateral symmetry and quality comparable to standard coronal and sagittal matrices. **(A)** Representative horizontal adult mouse brain section and quantification of the thickness of the fiber tract in the cc of the ACA (indicated by red dotted lines), length across HIP (indicated by white dotted lines), and number of HIP nuclei in **A’** (marked with black crosses) and its contralateral ROI **A’’.** Data from 1 of 2 independent experiments are show. N = 6 mice. Measurements were obtained from a series of horizontal sections across 200 µm in the dorsoventral axis. **(B)** Representative coronal adult mouse brain section and quantification of the thickness of the fiber tract in the cc (red dotted lines), length across HIP (white dotted lines), and number of HIP nuclei in **B’** (black crosses) and its contralateral ROI **B’’**. Data from N = 3 mice. Measurements were obtained from non-consecutive coronal sections spanning the rostral-caudal axis of the HIP area shown in **(A). (C)** Representative sagittal adult mouse brain section showing an alternate orientation of the corresponding HIP region seen in the horizontal **(A)** and coronal **(B)** sections. ACA, anterior cingulate area; CB, cerebellum; cc, corpus callosum; CP, caudate putamen; CTX, cortex; HIP, hippocampus; MB, midbrain; STR, striatum; TH, thalamus. Scale bars = 1 mm in horizontal, coronal, and sagittal overview sections; 100 µm in all magnified ROIs. Data in **(A)** and **(B)** are presented as mean ± SEM. Wilcoxon signed-rank test was performed for all left-right paired comparisons. No significant difference was found.

### Potential applications of the transversal brain matrix

The 1–3-mm-thick horizontal brain block cut from our transversal brain matrix is compatible with thin sectioning using a cryostat or microtome for downstream histological and immunofluorescence analyses. The involvement of minimal tissue handling, like the use of existing standard matrices, makes this protocol amenable to anterograde and retrograde tracing of local and long-projecting horizontal neuronal tracts by endogenous or adeno-associated virus-delivered fluorescence reporters (Burkhalter et al., 2024). Our protocol can be enhanced by the application of advanced tissue clearing methods such as CLARITY (Tian et al., 2024; Tyson et al., 2015) as well as benefit studies on interhemispheric circuitries (Liu et al., 2021; Szczupak et al., 2021; Tian et al., 2024) and subcortical brain nuclei (Antonovaite et al., 2018; Cao et al., 2020). Our device can complement widely used horizontal whole-brain imaging modalities such as magnetic resonance imaging (MRI) (Chuang et al., 2011; Wang et al., 2020) and X-ray microtomography or micro-CT (Humbel et al., 2025; Mizutani et al., 2016; Mrzílková et al., 2019) by post-imaging integration of modern cell and molecular technologies to study whole-brain connectivity at single-cell level in the same specimen. Skilled manipulation of our transversal matrix and cutting blades under chilled conditions may allow precise extraction of bilateral tissue punches from freshly perfused mouse brains for region-specific downstream single-cell -omic analysis in healthy and disease brains. Finally, users of our transversal matrix can benefit from the wealth of 3D reconstruction methods and brain atlases relevant to their areas of research (Agarwal et al., 2018; MacKenzie-Graham et al., 2004; Piluso et al., 2024; Stæger et al., 2020; Xiong et al., 2018).

In conclusion, our new transversal mouse brain matrix produced using CAD and 3D printing was a fast, practical, and cheap solution that has accelerated the experimental pipeline of our neurobiological research. We hope the broader neuroscience community benefits from our publicly available CAD files by reproducing our device locally through institutional or external 3D-printing resources, adapting our design for different brain sizes, and expanding the scope of their neurological investigations.

## ACKNOWLEDGEMENTS

We thank the BU Animal Science Center for providing the euthanized animals. Brain matrices were 3D printed at the BU Engineering Product Innovation Center (EPIC). This work was supported by BU startup funding awarded to TLT.

## AUTHOR CONTRIBUTIONS

KF, ABL, and RT designed and refined the transversal mouse brain matrix. KF generated the G-code. AK, HA, MCJ, and ACK processed the mouse brains. HA, MCJ, and ACK designed the histological experiment and executed the analysis. KF, AK, ABL, HA, MCJ, and TLT prepared the figures. ABL recorded the videos. CCA, ABL, HA, and RT drafted the manuscript. TLT conceived and supervised the study and finalized the manuscript.

## SUPPLEMENTARY FILES

## Supplementary Methods

**Supplementary Video 1 Cavity Angle.mp4**

**Supplementary Video 2 Cavity Curvature.mp4**

**Supplementary Video 3 Cavity Dimensions.mp4**

**Supplementary Video 4 Section Customization.mp4**

**Supplementary Video 5 Different Blade Sizes.mp4**

