## Supplementary Methods for "Reproducible transversal mouse brain sections using low-cost 3D printable resin matrix"

### Instructions for CAD drawing using Onshape (https://bu.onshape.com/)

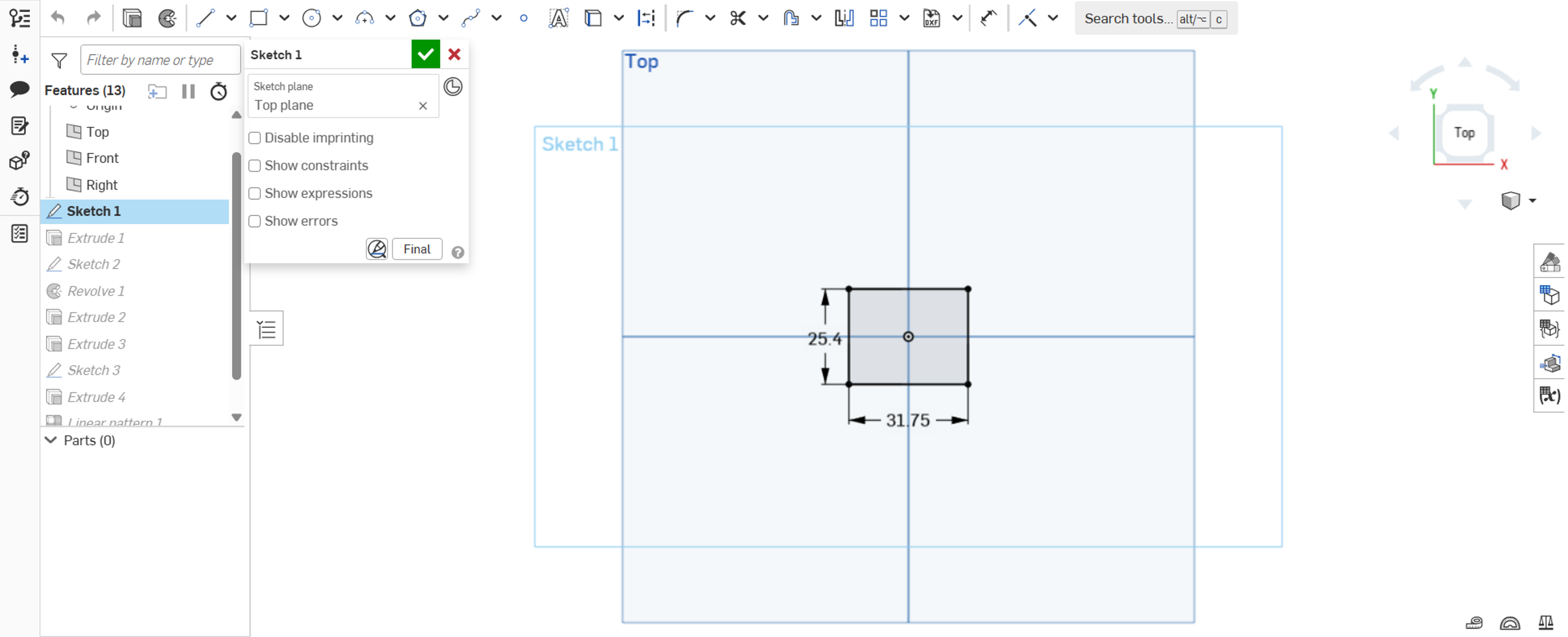


**Step 1.** Establish the base of the matrix (25.4 mm by 31.75 mm) using the Sketch and Center Point Rectangle tool.

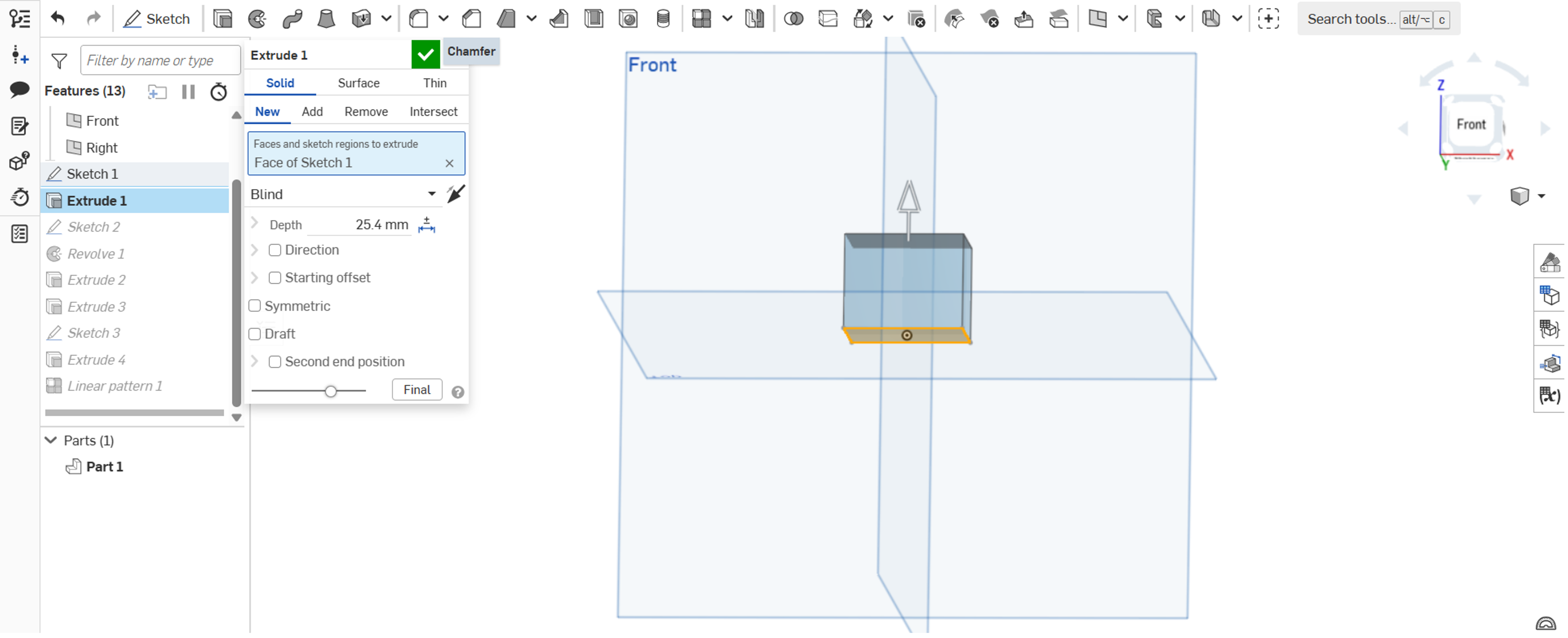


**Step 2.** Extrude the rectangle by 25.4 mm in the positive z-axis direction.

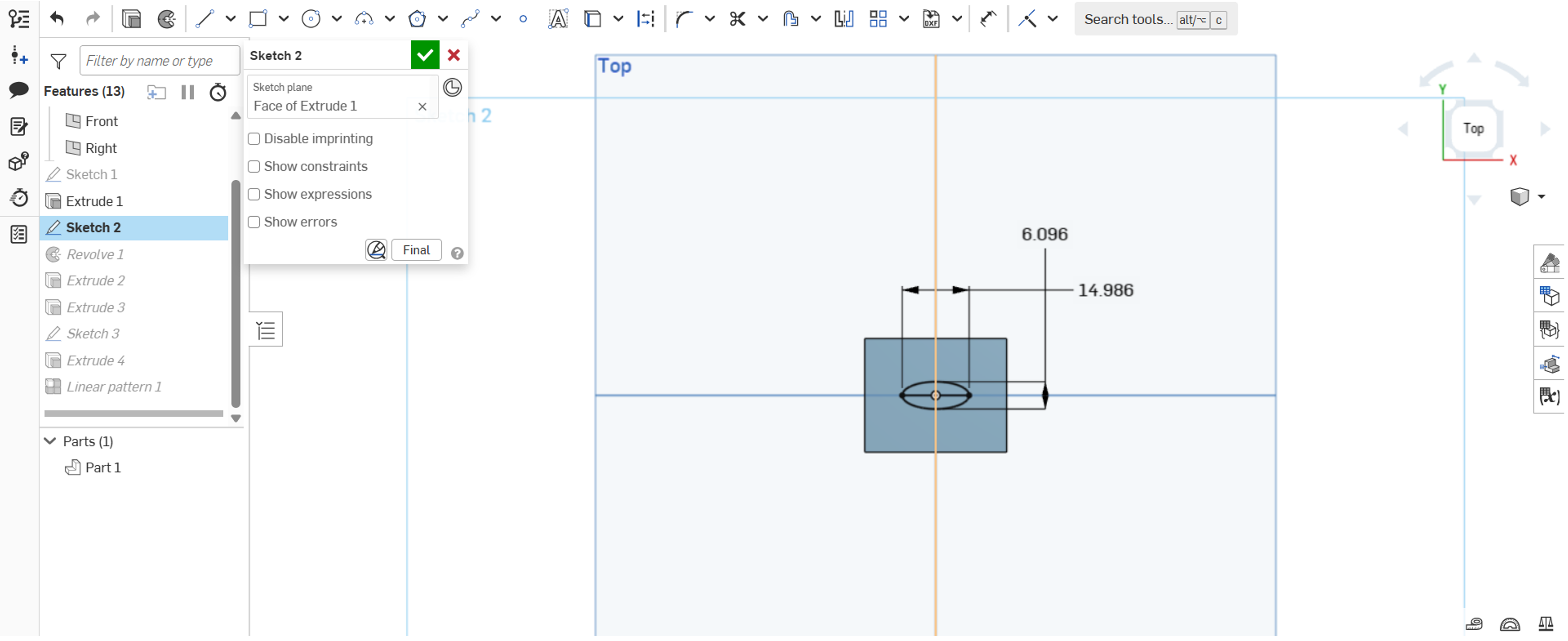


**Step 3.** Draw a 14.986 mm by 6.096 mm ellipse in the center of the device using the Sketch and the Center Point Rectangle tool to create the brain chamber.

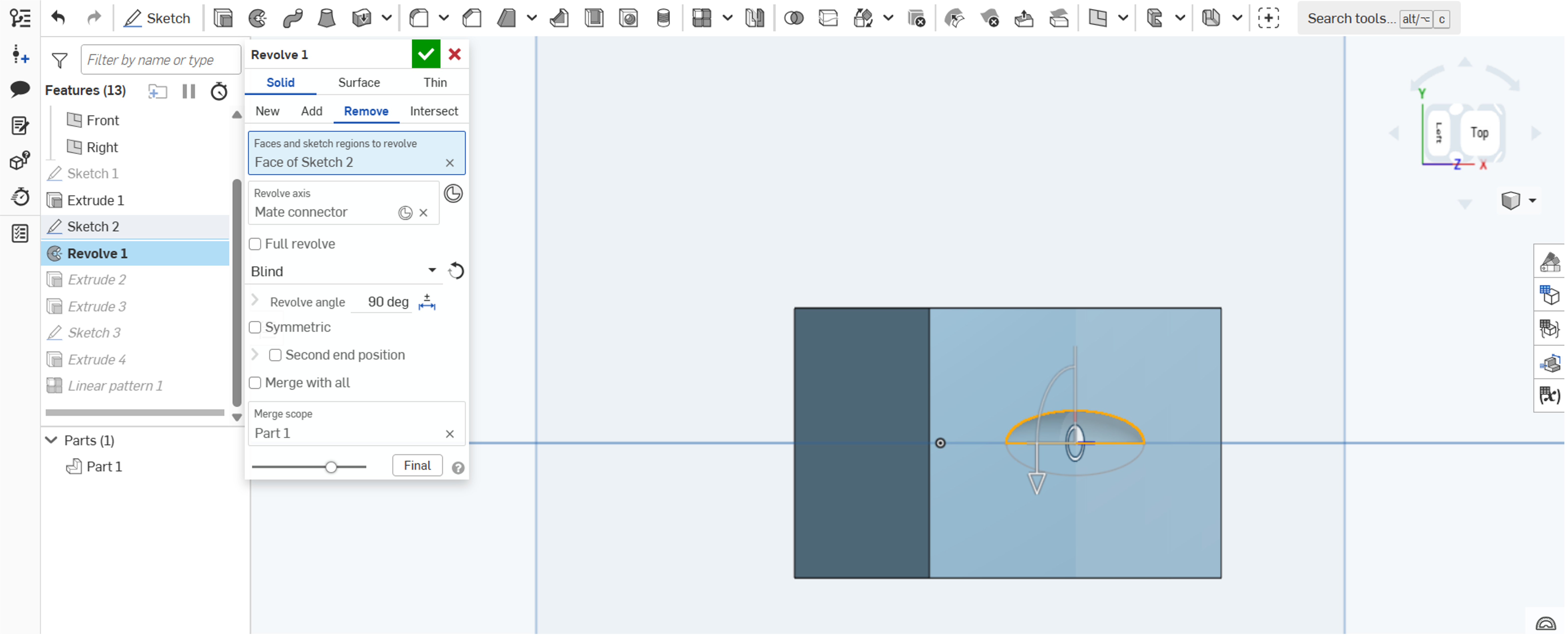


**Step 4.** Divide the ellipse into two symmetrical halves. Use the Revolve tool to rotate the top half of the ellipse 90° towards the negative z-axis.

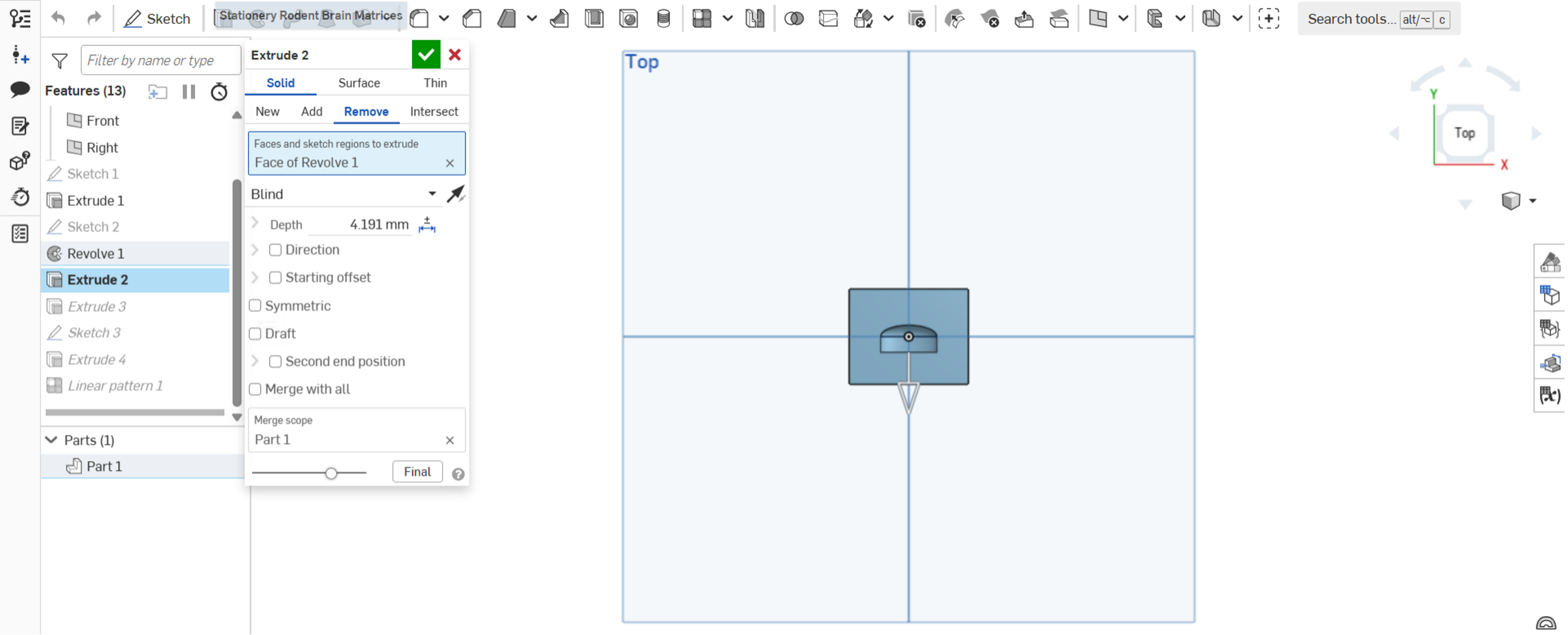


**Step 5.** Extrude the ventral-facing wall of the cavity laterally by 4.191 mm to fit the dorsal-ventral axis of an adult mouse brain.

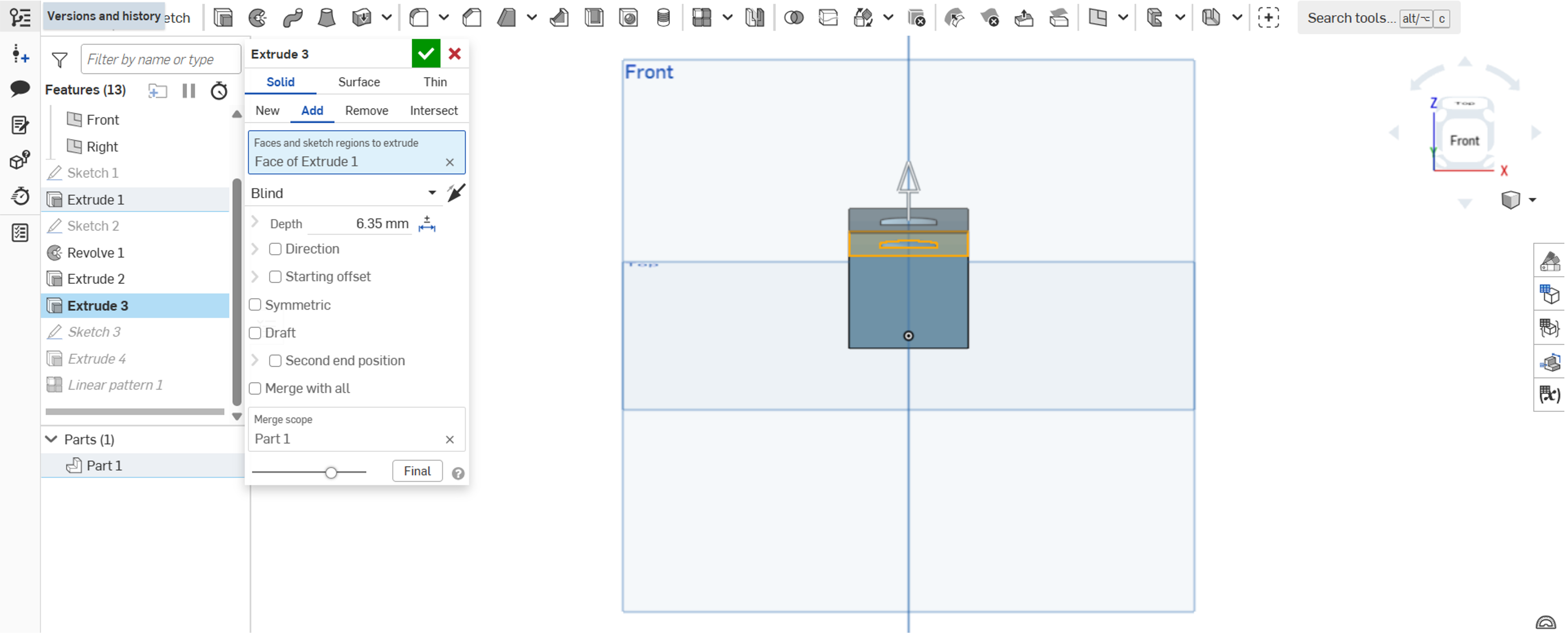


**Step 6.** Extrude the cavity by 6.350 mm to accommodate the horizontal width of the brain.

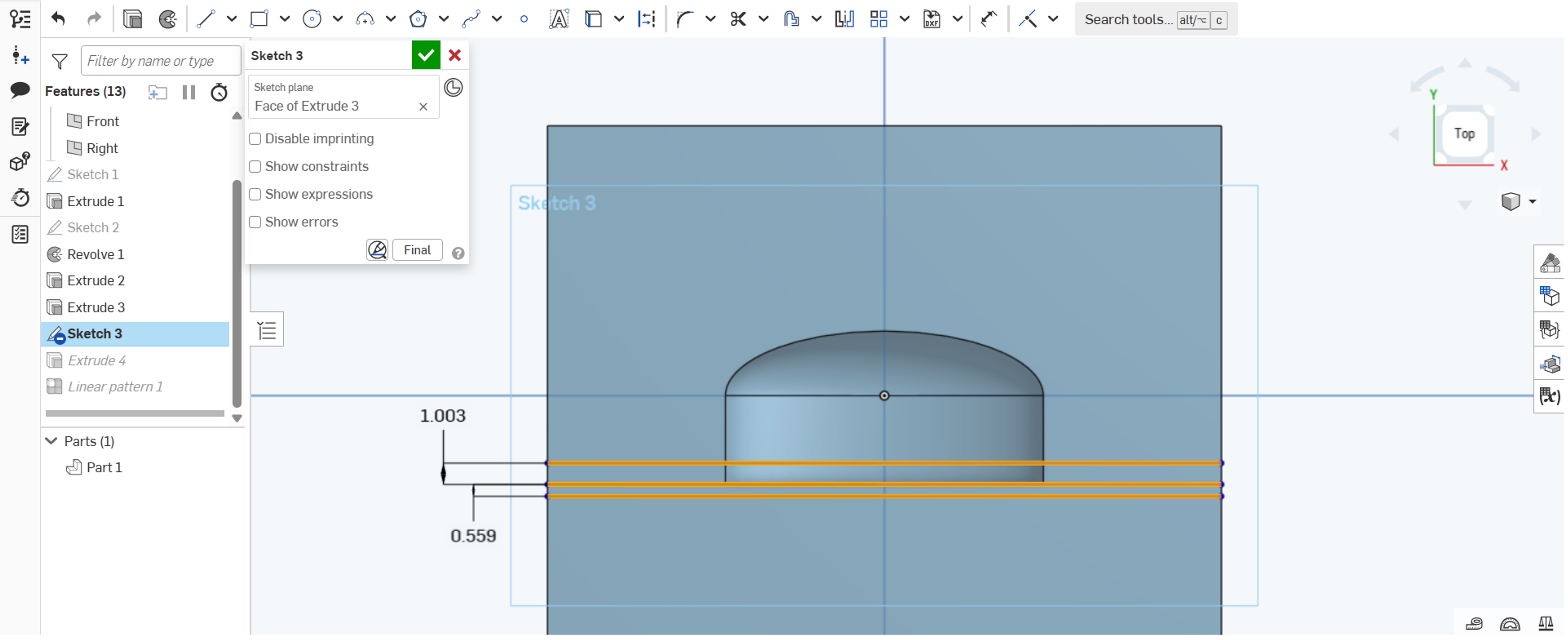


**Step 7.** Draw a 0.559-mm thick line using the Line tool to form the ventral-most slit of the brain matrix. Mark the location of the 1-mm wedge that alternate between slits.

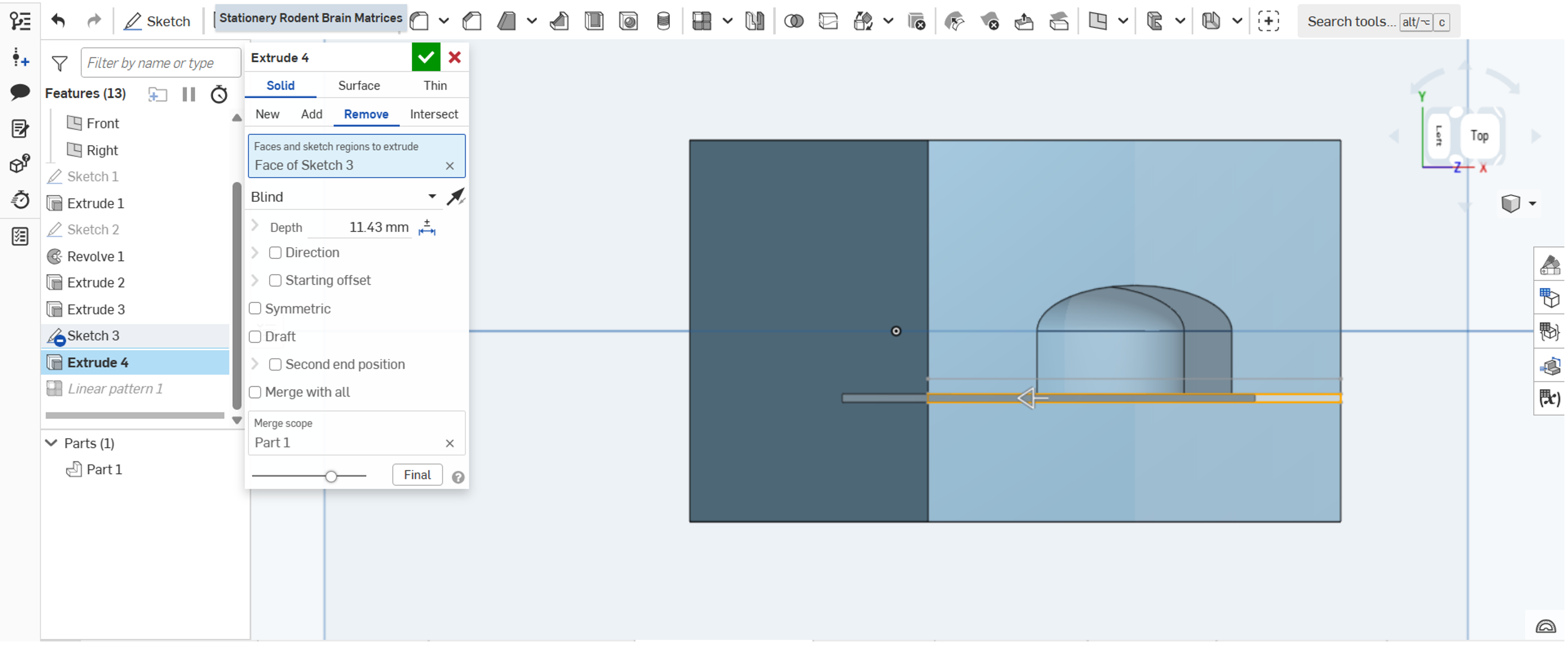


**Step 8.** Extrude the 0.559-mm slit 11.430 mm in the negative z-axis to establish the depth of the slit in the matrix.

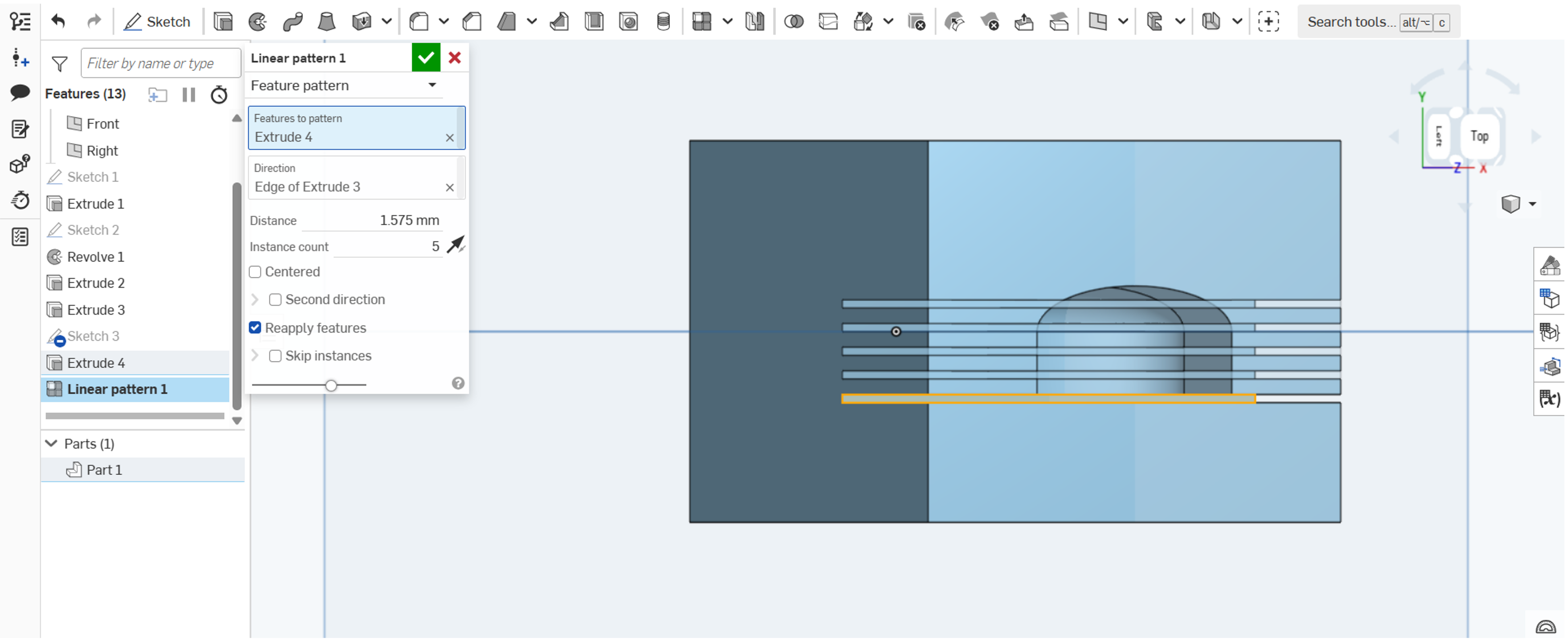


**Step 9.** Replicate the extruded slit using the Linear Pattern tool towards the opposite cavity wall (dorsal) to achieve a total of five slits.
